# Src-dependent DBL family members drive resistance to vemurafenib in human melanoma

**DOI:** 10.1101/561597

**Authors:** Charlotte R. Feddersen, Jacob L. Schillo, Afshin Varzavand, Hayley R. Vaughn, Lexy S. Wadsworth, Andrew P. Voigt, Eliot Y. Zhu, Brooke M. Jennings, Sarah A. Mullen, Jeremy Bobera, Jesse D. Riordan, Christopher S. Stipp, Adam J. Dupuy

## Abstract

The use of selective BRAF inhibitors (BRAFi) has produced remarkable outcomes for patients with advanced cutaneous melanoma harboring a *BRAF*^*V600E*^ mutation. Unfortunately, the majority of patients eventually develop drug-resistant disease. We employed a genetic screening approach to identify gain-of-function mechanisms of BRAFi resistance in two independent melanoma cell lines. Our screens identified both known and unappreciated drivers of BRAFi resistance, including multiple members of the DBL family. Mechanistic studies identified a DBL/Rac1/Pak signaling axis capable of driving resistance to both current and next-generation BRAF inhibitors. However, we show that the Src inhibitor, saracatinib, can block the DBL-driven resistance. Our work highlights the utility of our straightforward genetic screening method in identifying new drug combinations to combat acquired BRAFi resistance.

---

Melanoma is the deadliest form of skin cancer, with around 90,000 diagnoses of invasive disease and ∼10,000 deaths per year(1). Patients had few treatment options until the development of vemurafenib, a highly selective kinase inhibitor that specifically targets the *BRAF*^*V600E*^ mutant protein present in ∼50% of all melanoma cases(2). Initially, vemurafenib provided complete or partial response in over 50% of patients and increased progression-free survival (3). Unfortunately, most patients relapse once tumors acquire resistance to vemurafenib.

Genetic analysis of progression samples has identified resistance mechanisms, including amplification of *BRAF*^*V600E*^, expression of truncated *BRAF*^*V600E*^, and *RAS* mutation (4-6). However, these mechanisms explain only ∼60% of cases of BRAF inhibitor (BRAFi) resistance (5, 7, 8). Drug resistance can be delayed by combining vemurafenib with cobimetinib, a MEK inhibitor (MEKi), but most patients eventually develop progressive disease via resistance mechanisms that have not been well characterized (7). Thus, unexplained cases of resistance to MAPK inhibition (MAPKi) in human melanoma represent an important unmet clinical need.

Mechanisms of vemurafenib resistance have been studied in *BRAF*^*V600E*^ mutant human melanoma cell lines using genome-wide shRNA and CRISPR loss-of-function screens (9-11). Overall, these studies showed little overlap in candidate mechanisms. Two screens have been reported that attempted to identify drivers of vemurafenib resistance by high throughput over-expression of genes via lentiviral libraries (12, 13). Importantly, these screens failed to identify known mechanism of vemurafenib resistance (*e.g*. *BRAF*^*V600E*^ amplification or N-terminal truncation)(4, 6). These observations led us to develop a simple insertional mutagenesis screening approach using the *Sleeping Beauty* (SB) transposon system to identify novel drivers of vemurafenib resistance in an unbiased forward genetic screen.

The SB system is a well-established tool for developing mouse models of spontaneous cancer in which transposon-induced somatic mutations drive transformation (14). In this context, the SB system consists of two parts: a mutagenic transposon vector and the transposase enzyme. When introduced into the same cell, the transposase excises the transposon from a donor vector and integrates it a TA dinucleotide site in the host cell genome. In the context of some selective pressure (*e.g.* proliferation, drug treatment), cells with specific mutations conferring the ability to outcompete neighboring cells undergo clonal expansion, facilitating subsequent identification of these phenotype-driving mutations. This method has been used to select for specific phenotypes using *ex vivo* cell-based assays (15-17), as well as to drive the development of drug-resistant tumors in a engineered mouse models (18, 19). However, previous *ex vivo* approaches using human cells have been limited by the relative inefficiency of delivering both transposon and transposase vectors to cells. Moreover, prior studies have required the isolation of clonal cell populations to identify insertional mutations associated with the desired phenotype, a process that greatly reduces screen throughput. Collectively, these challenges have limited to broader application of SB mutagenesis in *ex vivo* screening approaches.

We present here the results of three SB mutagenesis drug resistance screens conducted in human *BRAF*^*V600E*^ mutant melanoma cells to identify novel drivers of resistance to either BRAFi treatment alone or BRAFi/MEKi combination treatment. Importantly, we detected recurrent N-terminal truncations of BRAF as a driver of BRAFi resistance, a mechanism previously associated with BRAFi resistance in human melanoma (6). We also identified *MCF2, VAV1, PDGFRB*, and N-terminally truncated RAF1 as drivers of BRAFi resistance. We experimentally verified the ability of candidates to drive drug resistance in independent melanoma cell lines, and analysis of transcriptome data clinical progression samples revealed that over-expression of our candidates is associated with BRAFi resistance in human patients. Finally, we elucidate a mechanism through which the DBL family members MCF2 and VAV1 act to drive drug resistance and show that this mechanism can be blocked with saracatinib, an inhibitor of the Src family. These findings demonstrate the utility of our genetic screening approach to identify clinically-relevant drivers of drug resistance, as well as the potential for discovering new therapeutic approaches to reverse or prevent its occurrence.

## RESULTS

### Sleeping Beauty mutagenesis drives drug resistance in human melanoma cells

We first engineered human melanoma cells (A375, SK-MEL-28) to express the hyperactive SB100x transposase (20). Cells stably expressing SB100x were subsequently transfected with either the mutagenic pT2-Onc3 transposon vector (21) or a control EGFP expression plasmid (**Fig. 1A**). Cells were then grown for 48 hours in standard culture conditions to allow the SB100x enzyme to integrate the mutagenic transposons into the genome of the transfected cells (**Fig. 1B**). Independent plates of mutagenized or control cells were pooled, and 1×10^6^ cells were seeded onto 10 cm plates. Cells were placed under drug selection 12-24 hours after plating, and genomic DNA was collected for analysis once drug-resistant colonies emerged.

**Figure 1.**
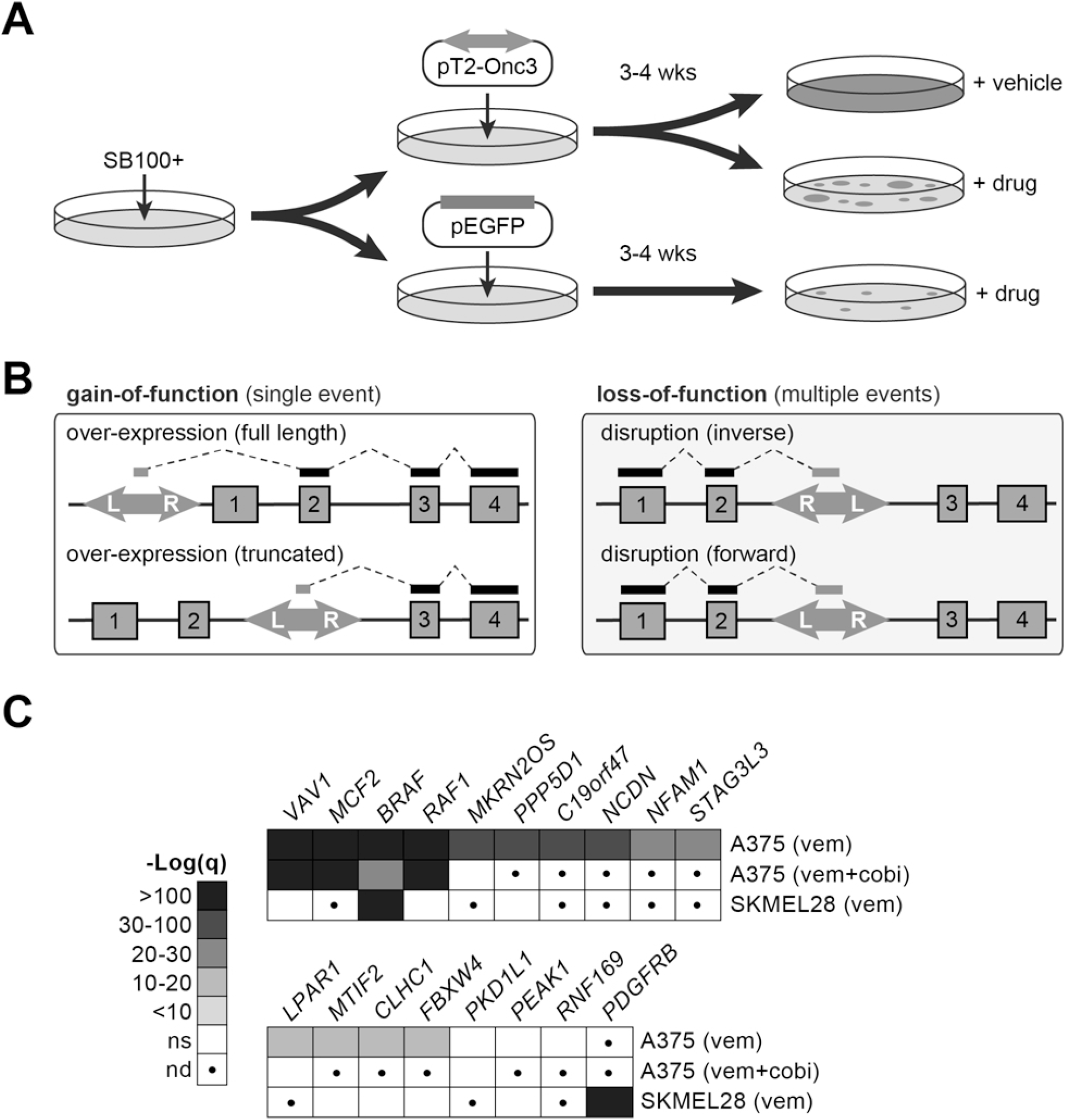
SB mutagenesis drives drug resistance in A375 through known and novel mechanisms. (**A**) A375 SB100x+ cells were transfected with the pT2-Onc3 transposon before undergoing drug treat with either vemurafenib (5 µm; n=75), vemurafenib plus cobimetinib (5µm and 5 nm respectively; n=20), or DMSO control (n=15). (**B**) The T2-Onc3 transposon can cause gain-of-function or loss-of-function mutations based on orientation of the transposon and location of insertion in the gene locus. (**C**) Drug resistance drivers were identified by profiling sites of transposon insertion in resistant cells to find genes that were recurrently over-expressed by transposon insertions. (**D**) Clustered transposon insertion sites within the *BRAF* locus are predicted to express an N-terminally truncated protein.

We generated a variety of mutagenized cell populations for our screen: vemurafenib-treated (5 µM), vemurafenib (5µM) combined with cobimetinib (5 nM), and control cells treated only with DMSO (*i.e.* vehicle). Vehicle-treated cells expanded rapidly and were collected ∼3 days after plating. Drug-resistant colonies emerged with varying kinetics on independent plates but generally appeared in ∼14 days in vemurafenib alone or ∼21 days in vemurafenib with cobimetinib. In both conditions, drug-resistant colonies appeared faster and in greater numbers than in non-mutagenized control cells (**Fig. S1A**), suggesting that transposon-induced mutations drive accelerated drug resistance.

Next, we determined if recurrent transposon-induced mutations could be identified in the resistant colonies. As picking individual colonies would limit the throughput of the approach, we instead harvested independent plates of cells as pooled populations, consistent with genome-wide shRNA and CRISPR protocols. We collected cell populations treated with vemurafenib [A375 (n=75), SKMEL28 (n=16)], vemurafenib plus cobimetinib [A375 (n=20], or vehicle [A375 (n=15), SKMEL28 (n=5)]. DNA fragments containing the transposon/genome junctions were amplified via ligation-mediated PCR (LM-PCR) and sequenced using the Illumina HiSeq platform as we have previously reported (22). Raw sequence reads were trimmed, mapped to the human reference genome (GRCh38), and filtered to remove rare insertion events.

The final dataset for each treatment group was analyzed using a modified gene-centric common insertion site (gCIS) analysis to identify genes with a higher rate of transposon insertion than predicted based on a random integration pattern (**Tables S1-S3**)(23). The screens identified a set of four genes in which over-expression is significantly associated with resistance to both vemurafenib and vemurafenib plus cobimetinib (**Fig. 1C**). Over-expression of thirteen additional genes was associated with resistance to vemurafenib alone. However, some of the differences between the two screens in A375 cell could be attributed to the differences in screen depth between the two drug conditions. Finally, differences in resistance mechanisms were also observed between the two cell lines. While over-expression of BRAF and RAF1 were common mechanisms in both A375 and SKMEL28, the DBL family of GEFs (MCF2) were unique to A375 while over-expression of PDGFRB was seen only in SKMEL28 cells (**Fig. 1C**, **Fig. S2**).

Closer inspection of the results revealed differences in the mechanism of resistance for individual genes. Transposon insertion within the promoter of first intron likely drives over-expression of full-length proteins in (**Fig. S2A,B**). The T2-Onc3 transposon is also capable of expressing truncated proteins (**Fig. S1B**). This mechanism appears to drive expression of N-terminal truncations of BRAF and RAF1 (**Fig. S2C,D**). Importantly, a truncation of the *BRAF*^*V600E*^ transcript in human melanoma has previously been shown create a similar N-terminally truncated protein associated with vemurafenib resistance in melanoma patients (6). Finally, the pattern of insertions in the *MCF2* locus suggest that over-expression of either full-length or truncated protein is associated with drug resistance (**Fig. S2E**). These mechanistic insights highlight the strength of transposon-based genetic screens to identify more complex mechanisms aside from simple knockdown or over-expression.

### Validation of candidate drug resistance drivers

We created vectors to mimic the transposon-induced expression of each gene. Each transgene was then stably expressed in A375 (**Fig. S3**). Drug resistance was evaluated by plating cells at low density in a 96-well culture format and serially measuring the relative viable cell number in each well over multiple days of culture in the presence of vemurafenib, vemurafenib plus cobimetinib, or vehicle (see Methods). Using this approach, we confirmed that over-expression of *VAV1, MCF2*, truncated *MCF2* (*MCF2*^Δ*N*^), truncated *BRAF* (*BRAF*^*V600E*^^Δ*N*^), or truncated *RAF1* (*RAF1*^Δ*N*^) significantly increases resistance to vemurafenib and vemurafenib plus cobimetinib (**Fig. 2A**). *BRAF*^*V600E*^^Δ*N*^ and *RAF1*^Δ*N*^ were the strongest drivers of resistance, with little growth inhibition upon drug treatment. Interestingly, full-length *RAF1* did not confer resistance to vemurafenib, illustrating the necessity of the N-terminal truncation (**Fig. 2A**). We also over-expressed *RAC1*^*P29S*^, a hotspot mutation known to drive BRAF inhibitor resistance (24), to assess the relative strength of our novel resistance drivers. *RAC1*^*P29S*^ performed similarly to VAV1 and MCF2 in our assay (**Fig. 2A**). Notably, none of the candidates identified in our screen (VAV1, MCF2, MCF2^ΔN^, BRAF^V600E^^ΔN^, RAF1^ΔN^) increased proliferation of A375 cells in the absence of drug. However, over-expression of RAC1, RAC1^P29S^, and full-length RAF1 did increase the growth rate significantly (not shown).

**Figure 2.**
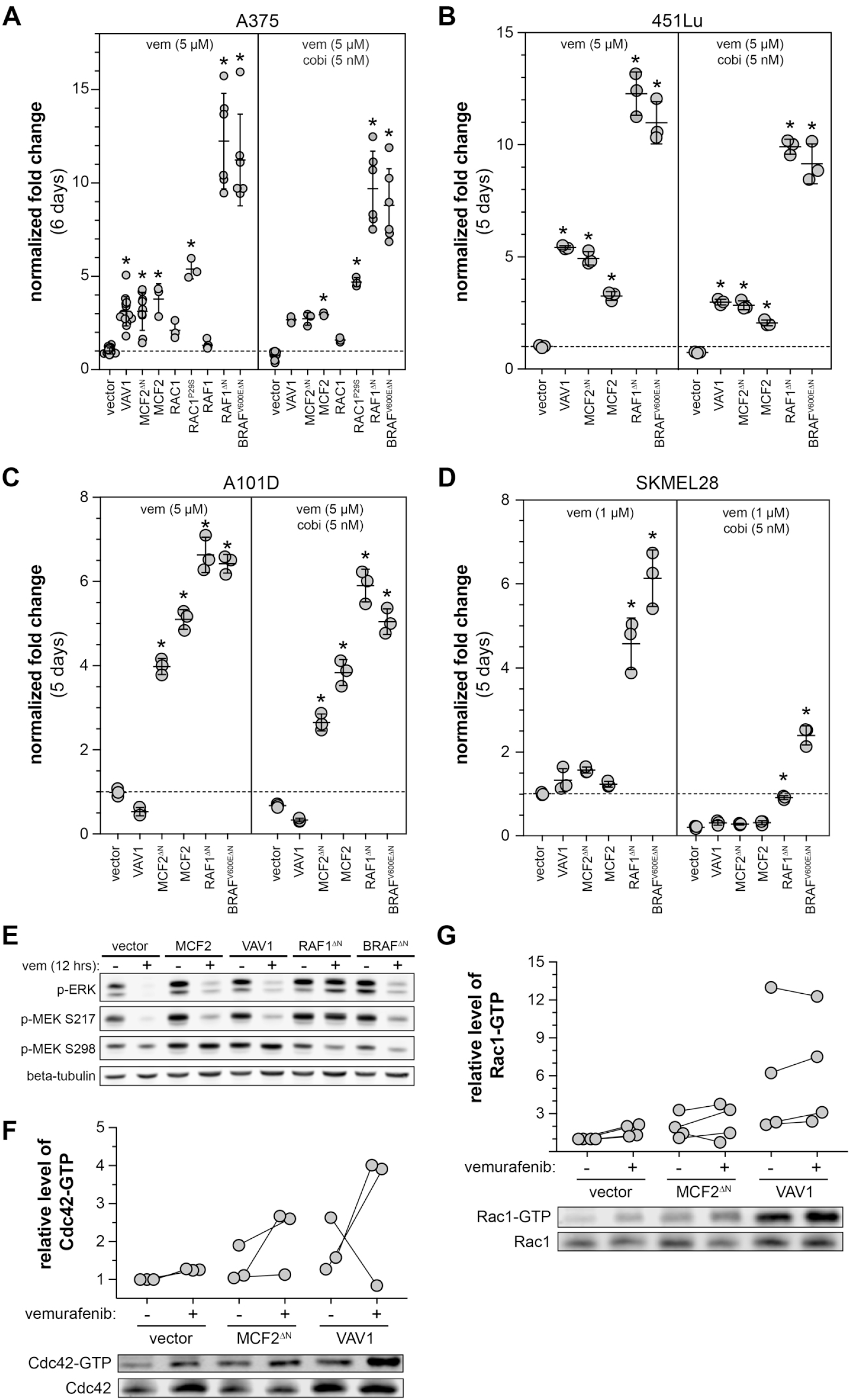
Validation of candidate drug resistance drivers in a panel of human melanoma cell lines. Growth of various engineered cell lines was assessed via CellTiter-Blue (Promega). The ability of candidates to increase resistance to vemurafenib and vemurafenib plush cobimetinib varied in (**A**) A375, (**B**) 451Lu, (**C**) A101D, (**D**) SKMEL28. [*corrected p-value ≤ 0.05, fold-change is relative to vector cells treated with 5 µm vemurafenib for each independent assay]. (**E**) A brief 12-hour vemurafenib treatment shows distinct patterns in MEK phosphorylation between the DBL- and RAF-driven mechanisms of resistance. Over-expression of either MCF2^ΔN^ or VAV1 increases the level of Cdc42-GTP (**F**) and Rac1-GTP (**G**).

### Performance of novel MAPKi resistance drivers in independent melanoma cell lines

Demonstration that our candidates confer drug resistance in A375 cells supports the validity of our genetic screen approach. To determine if the same mechanisms provide resistance in other MAPKi-sensitive *BRAF*^*V600E*^ mutant human melanoma cell lines, we generated populations of 451Lu, A101D, and SKMEL28 stably expressing each candidate gene and subjected them to the same 96-well growth assay for A375 cells. We determined that 451Lu had a similar resistance profile, with all of the candidates providing resistance to both vemurafenib and vemurafenib plus cobimetinib (**Fig. 2B**). All candidates were able to drive MAPKi resistance in A101D, with the exception of *VAV1* (**Fig. 2C**). Consistent with the findings from our genetic screen, only truncated BRAF and RAF1 were able to drive MAPKi resistance in SKMEL28, although the degree of resistance was much weaker than in A375 or 451Lu (**Fig. 2D**).

### DBL family members VAV1 and MCF2 drive MAPKi resistance through RAC1

While BRAF truncation is a well-established mechanism of MAPKi resistance, the mechanism of *VAV1* and *MCF2* driven resistance has not previously been investigated. Both VAV1 and MCF2 are members of the DBL family of guanine exchange factors (GEFs) and function by activating members of the Rho family of small GTPases(25). We evaluated MAPK signaling in the presence and absence of vemurafenib since MAPK reactivation is frequently observed in vemurafenib resistant tumors (26). Vemurafenib abolished phosphorylated ERK in vector control A375 cells (**Fig. 2E**). In contrast, phosphorylated ERK was partially restored in cells expressing MCF2, VAV1, and BRAF^ΔN^, and completely restored in RAF1^ΔN^. Next, we examined MEK phosphorylation upstream of ERK in cells expressing each candidate. As expected, expression of either truncated RAF1 or BRAF^V600E^ partially or completely restored phosphorylation at the RAF-controlled serine 217 on MEK in cells exposed to vemurafenib (**Fig. 2E**). In contrast, over-expression of VAV1 or MCF2 promoted maintenance of phosphorylation of serine 298 on MEK, an indication of elevated Pak activity in these cells (**Fig. 2E**). This observation is consistent with the finding that increased Pak signaling can drive acquired resistance to MAPK inhibitors in melanoma(27). Taken together, the dichotomous activation of MEK suggests a mechanistic distinction between resistance driven by RAF truncation versus DBL over-expression.

We next investigated how VAV1 and MCF2 work to reestablish ERK signaling in the presence of MAPKi. Both VAV1 and MCF2 were originally identified as oncogenes that act as guanine exchange factors for the Rho family(28, 29). Although the precise Rho family targets for these proteins are not entirely clear, it is generally accepted that VAV1 and MCF2 have activity for Rho, Rac1, and Cdc42(30). We verified that over-expression of VAV1 and MCF2^ΔN^ increases the levels of GTP-bound Cdc42 (**Fig. 2F**) and Rac1 (**Fig. 2G**), particularly in the presence of vemurafenib.

Over-expression of VAV1 and MCF2 could drive vemurafenib resistance via a Rac1/CdcD42/Pak pathway and/or by signaling via Rho through Rho-associated kinase (Rock)(**Fig. 3A**). To distinguish between these mechanisms, we assessed if inhibition of either Rock or Pak alters vemurafenib resistance driven by MCF2 or VAV1. First, we identified drug concentrations that had a minimal impact on cell growth in the absence of vemurafenib (**Fig. S4A,B**). Inhibition of Rock with Fasudil (10 µM) had variable effects on vemurafenib resistance across cell populations. However, vemurafenib resistance was consistently reduced by the addition of the Pak inhibitor FRAX-486 (50 nM) (**Fig. 3B**). Inhibition of Pak also prevented the emergence of spontaneous vemurafenib resistance in long-term A375 cultures (**Fig. 3C**), while Rock inhibition appeared to accelerate spontaneous resistance (**Fig. 3D**). Finally, knockdown of Rac1 in parental A375 showed synergistic cell killing in long-term culture with vemurafenib (**Fig. 3E**) but not vehicle (*i.e.* DMSO)(**Fig. S4D**). Conversely, knockdown of RhoA or RhoC did show any significant effects in parental A375 under standard culture conditions (**Fig. 3F**, **Fig. S4E**).

**Figure 3.**
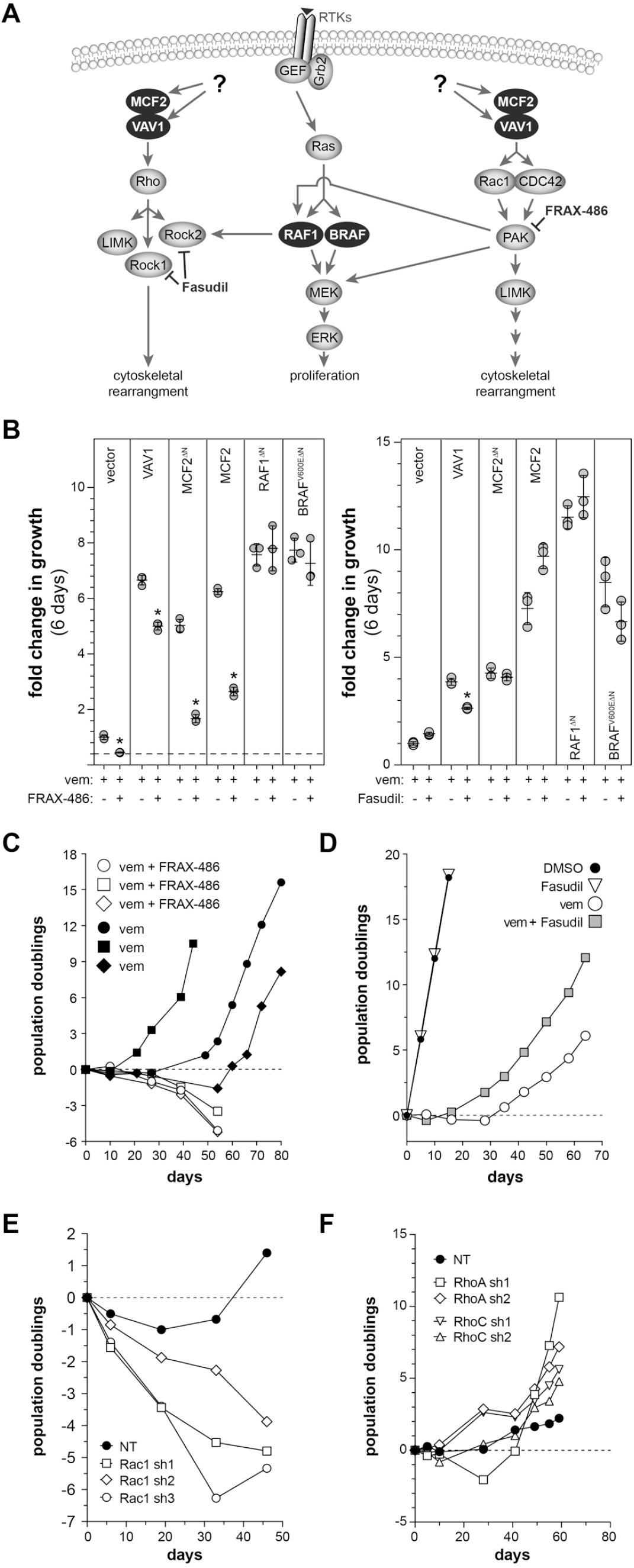
Evaluation of DBL GEF downstream signaling mechanism in A375 cells. (**A**) DBL family members have exchange activity for members of the Rho and Rac/Cdc42 family, each having distinct signaling mechanisms. (**B**) The addition of the Pak inhibitor FRAX-486 (50 nM) is able to reduce vemurafenib resistance driven by MCF2 and VAV1. However, the Rock inhibitor Fasudil did not show a consistent effect [*adjusted p < 0.001 relative to vemurafenib alone]. (**C**) Long-term treatment of A375 cells with FRAX-486 is capable of converting vemurafenib-induced cytostasis to cell killing, while long-term treatment with Fasudil (**D**) shows a trend of enhanced growth in vemurafenib. (**E**) Knockdown of RAC1 using three independent shRNAs converts vemurafenib-induced cytostasis to cell killing in long-term cultures of A375, while knockdown of either RhoA or RhoC does not impact cell growth with long-term vemurafenib treatment (**F**).

### The role of novel vemurafenib resistance drivers in spontaneous melanoma progression

While N-terminal truncations of BRAF have previously been associated with vemurafenib resistance in patients (6), the other candidate resistance drivers we identified have not. We sought to determine if alterations in these candidates would arise spontaneously to provide vemurafenib resistance in the absence of experimentally-induced mutagenesis. First, we generated A375 cells with spontaneously acquired resistance after long-term culture in 3 µM vemurafenib. Two phenotypically distinct populations of resistant cells were derived from this process. The first vemurafenib-resistant cells that grew out of these cultures were derived from colonies showing a compact morphology. We isolated nine such colonies from eight different populations of vemurafenib-treated A375 cells. We also isolated three independent populations of the second class of vemurafenib-resistant A375 cells, which grew more diffusely and slowly. Importantly, cell populations of this second class do not generate the more rapidly growing colonies of vemurafenib resistant cells.

We first looked for evidence of BRAF^V600E^ alterations in the nine colonies and three populations of vemurafenib-resistant A375 cells by performing western blotting using an antibody specific to BRAF^V600E^ isoform (**Fig. S5A**). No alterations were detected in the vemurafenib-resistant populations (VRP1-3), but six of the nine vemurafenib-resistant clones showed evidence of BRAF^V600E^ protein alteration (**Fig. S5A**). Two of the six clones express a truncated form of BRAF^V600E^ that is ∼40 kD, consistent with truncated forms of BRAF^V600E^ known to drive vemurafenib resistance in patients (6). We confirmed that this truncation is caused by an intragenic deletion of BRAF exons 2 through 10 (data not shown), leading to the expression of an altered form of the BRAF^V600E^ transcript that splices directly from exon 1 into exon 11. An additional four vemurafenib-resistant clones appear to express a BRAF^V600E^ fusion that is ∼120 kD in size (**Fig. S5A**). Consistent with the presence of BRAF alterations, the level of pMEK-S217 (RAF phosphorylation site) was elevated in all but one of the vemurafenib-resistant clones, even in the presence of vemurafenib (**Fig. S5B**). A similar approach did not provide evidence of RAF1 alterations in vemurafenib-resistant clones and populations (data not shown).

Next, we performed western blotting to determine if either MCF2 or VAV1 expression is altered in the A375 cells with acquired spontaneous resistance to vemurafenib. While no changes in MCF2 expression were detected, VAV1 protein expression is significantly higher in all three vemurafenib-resistant cell populations (VRP1-3) (**Fig. S5B**). Levels of pMEK-S298 were also elevated in these populations, consistent with our findings that VAV1 drives vemurafenib resistance via Rac1/Pak signaling (**Fig. S5B**). The increased levels of VAV1 protein were not accompanied by increased mRNA expression (data not shown), suggesting that the increased protein level is achieved through post-transcriptional regulation (*e.g.* increased protein stability). These results provide corroborating evidence that VAV1 is involved in spontaneous vemurafenib resistance.

Finally, we used several independent approaches to evaluate the novel candidate vemurafenib resistance drivers (RAF1, MCF2, VAV1) in human melanoma samples. First, we analyzed 159 *BRAF*^*V600E*^-mutant melanoma samples present in the Cancer Genome Atlas cutaneous melanoma project (TCGA-SKCM). Although BRAFi response data are not available for these samples, we evaluated the expression levels of candidates identified in our vemurafenib resistance screen that are predicted to drive resistance when over-expressed (**Table S1**). Expression of VAV1 initially appeared to be elevated in a portion of melanomas in the TCGA-SKCM panel. However, it has been shown that VAV1 is highly expressed in lymphocytes. Indeed, further analysis showed that VAV1 expression correlated strongly with T-cell markers, suggesting that the majority of VAV1 expression in the TCGA-SKCM biopsies was contributed by infiltrating lymphocytes. Therefore, we were unable to evaluate VAV1 expression using transcriptome sequencing data derived from bulk analysis of melanoma biopsies. However, each of the remaining genes showed over-expression (z-score ≥ 2) in a subset of TCGA-SKCM samples. Overall, 56 samples (∼37%) exhibit over-expression of one or more genes from the set (**Fig. 4A**).

**Figure 4.**
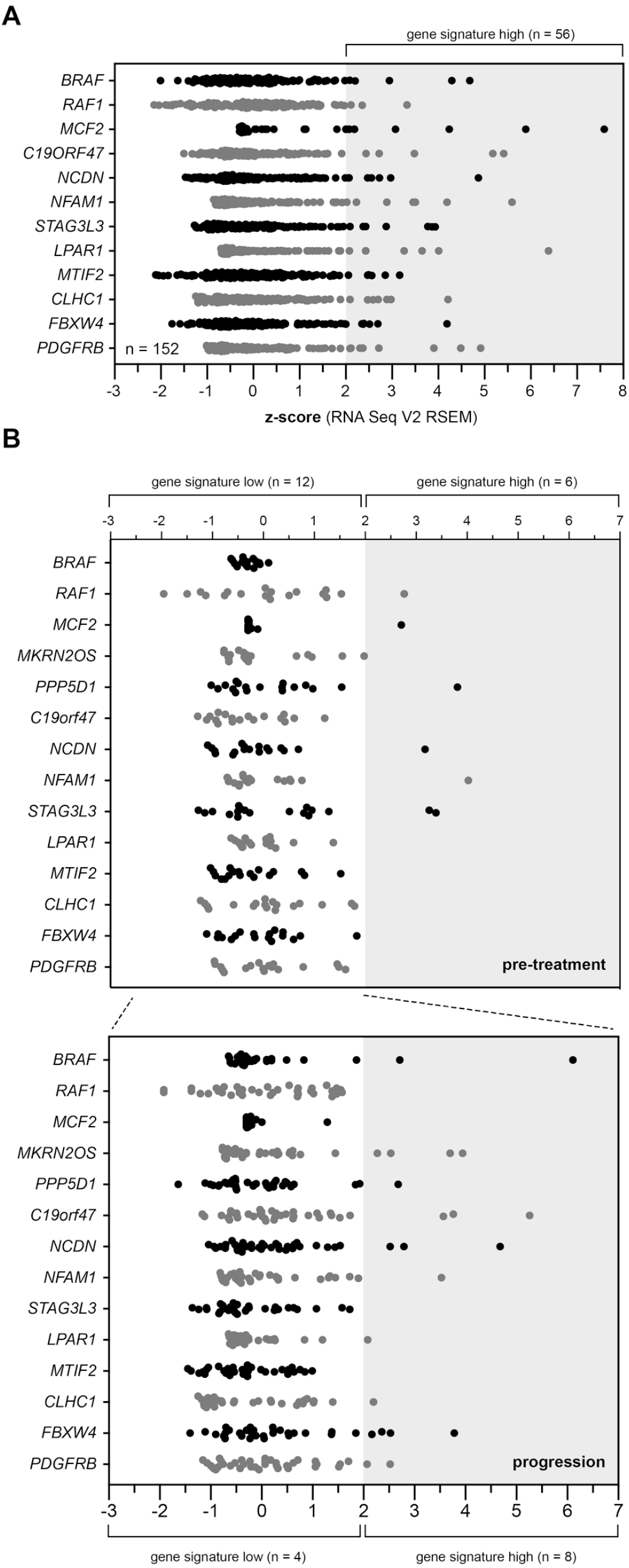
Role of candidate drug resistance drivers in human melanoma patients. (**A**) Evaluation of the top vemurafenib resistance drivers in primary cutaneous melanoma samples from the TCGA-SKCM project. Over 37% of patient samples show increased expression of at least one of the candidate resistance drivers. (**B**) Expression of the candidate resistance drivers in an independent RNA-seq data set obtained from a cohort of 18 patients, each with a diagnostic sample (*i.e.* pre-treatment) matched with a second sample taken after progression on BRAFi therapy (31). (above) Six of the 18 patients had elevated expression of at least one resistance driver in the pre-treatment sample. An additional eight patients show over-expression of at least one candidate resistance driver in the progression samples.

Unfortunately, the majority of samples in the TCGA-SKCM panel lack sufficient patient treatment histories to allow correlation of gene expression with response to MAPKi treatment. Therefore, we evaluated an RNA-seq data set obtained from a collection of primary melanomas with matched progression samples(31). Hugo *et al*. obtained RNA-seq data on pre-treatment melanoma samples taken from eighteen patients along with matched biopsies obtained after patients had progressed during treatment with a BRAFi alone or a BRAFi/MEKi combination.

We evaluated expression of the vemurafenib resistance gene set using this RNA-seq data to identify patterns in expression that correlate with treatment response. Pre-treatment tumor samples showed an expression pattern similar to that observed in the TCGA-SKCM panel with 6 of 18 (∼33%) patient samples showing over-expression of at least one of the resistance driver genes (**Fig. 4B**). We then examined expression of the gene set in progression samples (n=33) taken from patients whose primary tumor sample did not initially exhibit over-expression. Interestingly, eight of twelve patients (∼66%) appeared to acquire over-expression of at least one resistance driver (**Fig. 4B**).

### Src inhibition blocks MAPKi resistance driven by Rac signaling

Our forward genetic screen identified several members of the DBL family of guanine nucleotide exchange factors (MCF2, MCF2L, VAV1, VAV2) as novel drivers of vemurafenib resistance (**Fig. 1C**). Unfortunately, there are no drugs available that can inhibit the activity of GEFs or their small GTPase targets. However, the activity of the DBL family is regulated by a variety of upstream signals. For instance, both VAV1 and MCF2 can be activated by Src-mediated phosphorylation(32, 33). Interestingly, several prior publications have shown that the Src inhibitor saracatinib exhibits synergism with vemurafenib, although this synergism was not attributed Src’s role in regulating VAV1 or MCF2(34, 35).

We hypothesized that inhibiting Src using saracatinib would block vemurafenib resistance driven by MCF2 and VAV1 over-expression. We performed a short-term growth assay in A375 cells engineered to express each resistance driver. As we had seen previously, expression of MCF2, VAV1, RAC1^P29S^, RAF1^ΔN^, and BRAF^ΔN^ all drove growth of A375 cells in vemurafenib and vemurafenib combined with cobimetinib (**Fig. 5A**). As predicted, the addition of saracatinib to vemurafenib inhibited the growth of cells over-expressing MCF2 and VAV1. Surprisingly, saracatinib was also able to block vemurafenib resistance driven by RAC1^P29S^ expression, suggesting that the P29S activation mechanism may still depend on Src-dependent GEF activity. In all of these cases, the combination of vemurafenib and saracatinib not only blocked growth but caused cell death over the course of the 12-day assay (**Fig. 5A**). Although vemurafenib resistance driven by truncated BRAF^V600E^ and RAF1 was modestly reduced by the addition of saracatinib, these cells were still able to grow in the presence of both drugs (**Fig. 5A**).

**Figure 5.**
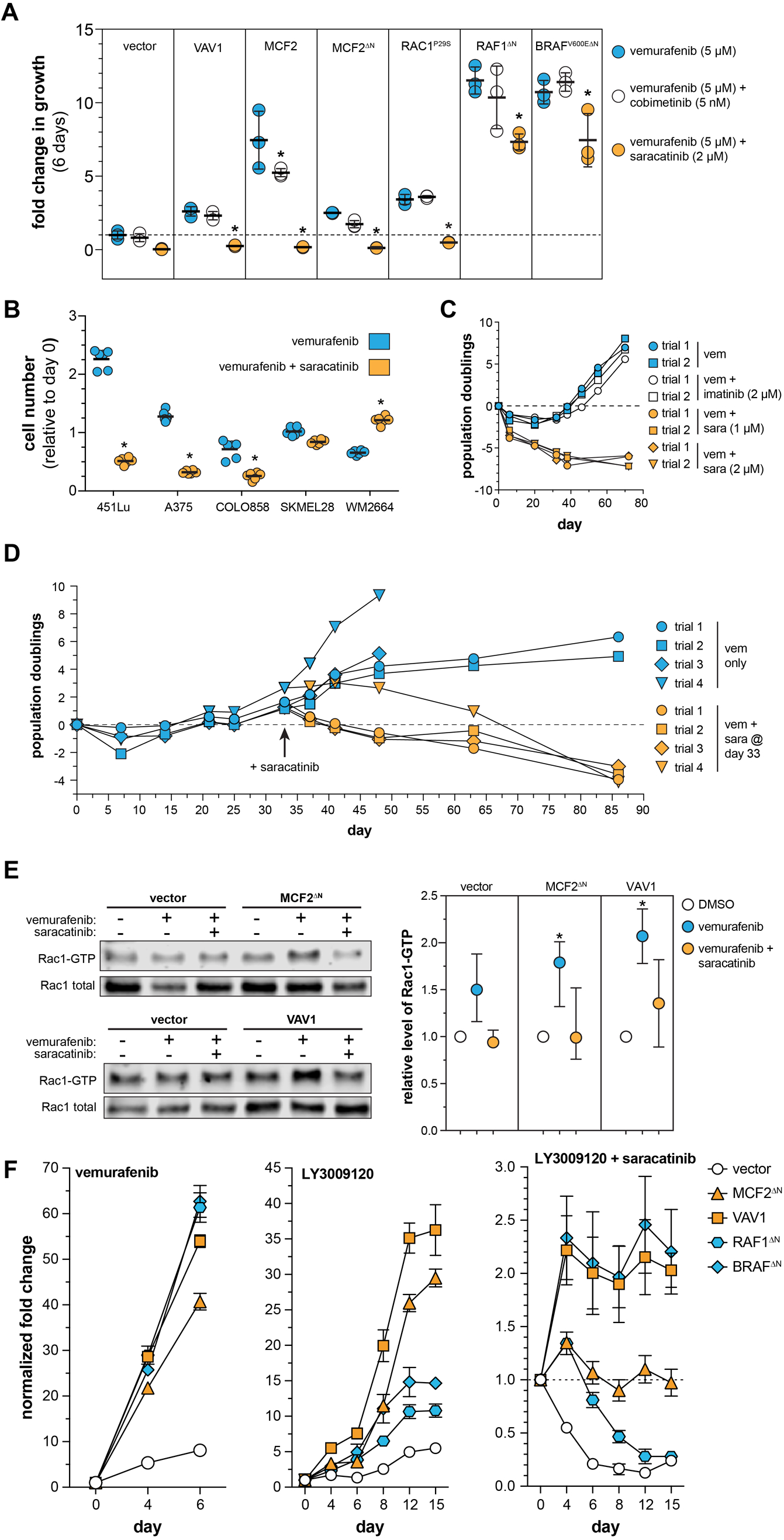
A Src inhibitor, saracatinib, shows synergistic cytotoxicity in combination with a next generation Raf inhibitor. (**A**) The resistance driven by MCF2 and VAV1 can be overcome with the addition of saracatinib. However, saracatinib cannot block resistance driven by the expression of truncated BRAF^V600E^ or RAF1 [*adjusted p-value < 0.05 relative to vemurafenib alone]. (**B**) The addition of saracatinib does not uniformly increase vemurafenib response in a panel of *BRAF*^*V600E*^ mutant human melanoma cell lines [*adjusted p-value < 0.0001]. (**C**) Saracatinib, but not imatinib, is able to induce cytotoxicity with vemurafenib in A375 cells. (**D**) The addition of saracatinib is able to induce cytotoxicity in cells that have acquired spontaneous resistance to vemurafenib. (**E**) Saracatinib treatment reverses the increase in Rac1 activation observed in A375 cells expressing MCF2 and VAV1. (**F**) The resistance candidates perform differently in response to treatment with LY3009120, a next generation Raf inhibitor. Importantly, the combination of LY3009120 and saracatinib is cytotoxic to all cell populations (note independent y-axes).

We next determined if the addition of saracatinib increases vemurafenib sensitivity across a panel of *BRAF*^*V600E*^ mutant melanoma cell lines. Saracatinib exhibited synergistic cell killing in short-term growth assays with vemurafenib in three cell lines (A375, 451Lu, COLO858) that was not observed in two additional cell lines (SKMEL28, WM2664) (**Fig. 5B**). Prior work has shown that saracatinib can inhibit the ABL kinases, albeit with lower activity(36). We performed long-term growth assays using A375 cells to determine if ABL kinase inhibition using imatinib had a similar effect as saracatinib. Two different concentrations of saracatinib were able to block the emergence of A375 cells with spontaneous vemurafenib resistance while imatinib treatment did not impact the acquisition of spontaneous resistance (**Fig. 5C**). Moreover, saracatinib was also able to kill A375 cells after spontaneous vemurafenib resistance developed (**Fig. 5D**). Finally, the addition of saracatinib reverses the activation of Rac1 observed in A375 cells over-expressing MCF2^ΔN^ and VAV1, consistent with Src acting upstream as an activator of DBL GEF activity (**Fig. 5E**).

As previously mentioned, N-terminal truncations of BRAF have been shown to drive resistance to vemurafenib in patients(6). Subsequent work has shown that BRAF truncations and fusions can drive resistance by functioning as constitutively-active dimers, which cannot be blocked by vemurafenib(6, 37, 38). However, next generation BRAF inhibitors (*e.g*. LY3009120) have been developed that are capable of inhibiting Raf dimers in addition to monomeric BRAF^V600E^ (37, 38). These dimer-blocking compounds are active against RAS mutant cells because they are able to inhibit RAS-dependent BRAF dimers, which cannot be blocked by vemurafenib(37). Nevertheless, cells with intrinsic resistance to LY3009120 have been reported(37), suggesting that not all BRAFi resistance mechanisms act by enforcing BRAF dimerization.

We tested the ability of the dimer-blocking drug LY3009120 to inhibit growth of A375 cells expressing each of the resistance drivers identified by our screen. As expected, LY3009120 significantly reduced the growth of cells expressing either truncated BRAF^V600E^ or RAF1 (**Fig. 5F**). However, expression of MCF2^ΔN^ and VAV1 were still able to drive proliferation in the presence of LY3009120, suggesting that the Rac-driven resistance mechanism may not rely on RAF dimerization. Nevertheless, the combination of LY3009120 and saracatinib was able to induce cytotoxicity in all cell populations, suggesting that this drug combination can thwart both resistance mechanisms identified by our genetic screen.

Each drug resistance driver was then tested in A375 cells using a panel of MAPK inhibitors including four RAF inhibitors, three MEK inhibitors, and an ERK inhibitor (**Fig. 6A**). These experiments demonstrated several trends. First, none of the currently approved RAF (vemurafenib, dabrafenib, encorafenib) or MEK inhibitors (cobimetinib, trametinib, binimetinib) were able to control the growth driven by the resistance drivers as mono- or combination therapies (*i.e.* RAFi + MEKi). The ERK inhibitor ulixertinib was also unable to suppress the growth of cells over-expressing MCF2 and VAV1. However, while saracatinib (SRCi) alone had little effect on cell growth, the combination of RAFi with SRCi was much more effective. The combination of LY3009120 with saracatinib was the most effective drug combination (**Fig. 6A**). As previously observed, RAFi with SRCi was effective in suppressing the growth of 451Lu (**Fig. 6B**) but not A101D or SKMEL28 (**Fig. 6C,D**).

**Figure 6.**
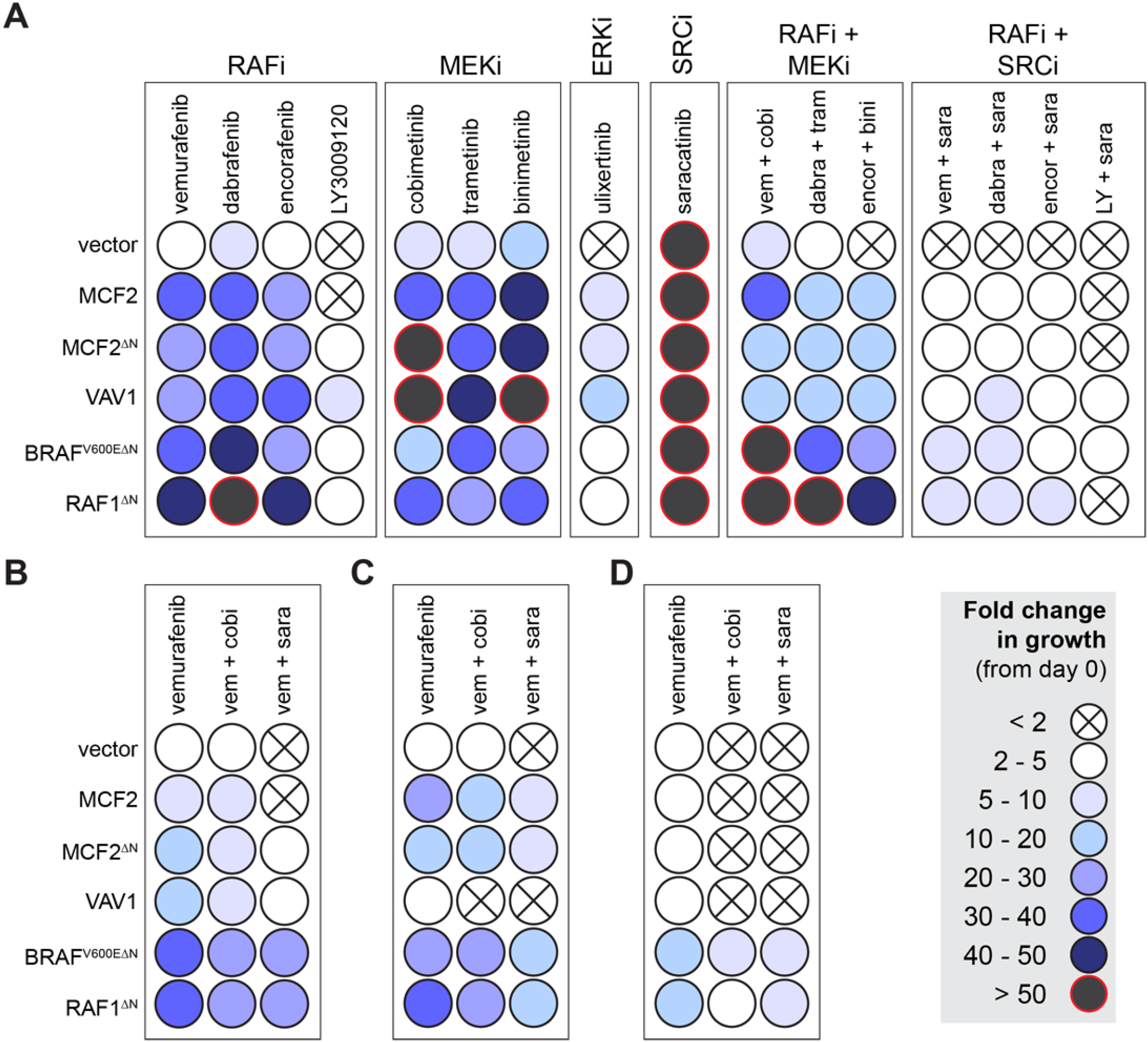
Performance of candidate resistance drivers across a panels of MAPK inhibitors. The ability of candidates to increase resistance to the indicated drugs is shown as a relative colorimetric endpoint (see legend). Each candidate was tested in the indicated drug combination in (**A**) A375), (**B**) 451Lu, (**C**) A101D, and (**D**) SKMEL28. Rows represent independent populations engineered to express the indicated candidate drug resistance driver shown at the left.

## DISCUSSION

We report here the results of a forward genetic screen designed to identify gain-of-function mutations that drive resistance to targeted kinase inhibitors (**Fig. 1A**). Our approach utilized a hyperactive form of the Sleeping Beauty transposase to increase the mutagenesis efficiency in cultured cells, allowing us to establish a simple screening method is easily replicated and applied in other contexts. We used this approach to identify novel drug resistance drivers in a variety of conditions. The screen results suggest that while drug resistance can be driven by common mechanisms (*e.g.* BRAF/RAF1 truncation), there are diverse mechanisms that are unique to specific drug combinations or cell lines (**Fig. 1C**). However, additional work is needed to validate the broader findings from our screens.

Prior work has already shown that N-terminal truncation of BRAF is associated with vemurafenib resistance in patients(6). We observed transposon clustering in the middle of the *BRAF* locus predicted to drive expression of a similar truncated isoform of BRAF (**Fig. S2C**), strongly supporting the relevance of our screening method. Furthermore, we observed a similar pattern of insertions in the *RAF1* locus (**Fig. S2D**), and we subsequently showed that expression of an N-terminal truncated isoform of RAF1 can drive drug resistance to a level comparable to that of truncated BRAF in a collection of vemurafenib-sensitive human melanoma cells (**Fig. 2**). This finding suggests that *RAF1* N-terminal truncation through intragenic deletion or gene fusion events could account for resistance in some human melanoma patients.

The other major finding from our genetic screens is that over-expression of the DBL-family guanine nucleotide exchange factors MCF2 and VAV1 can drive resistance to MAPK inhibition (**Fig. 6**). While the DBL family of GEFs act on both Rho and Rac proteins (25), we show that the DBL-driven resistance likely signals through Rac/Cdc42/Pak rather than Rho/Rock (**Fig. 3**). A prior study showed that gain-of-function mutations in *RAC1* are associated with vemurafenib resistance(24), and over-expression of the DBL GEFs is another mechanism through which RAC1 can be activated to drive resistance. To our knowledge, this is the first time the DBL family has been implicated in vemurafenib resistance.

All of the candidate resistance drivers we chose for further study significantly increased drug resistance in at least two independent sensitive human melanoma cell lines (**Fig. 2**). To further establish the relevancy of our candidates, we also evaluated the candidate drivers for a role in vemurafenib resistance by studying 12 independent clonal populations of A375 cells that had spontaneously acquired vemurafenib resistance, finding evidence of BRAF^V600E^ alterations (**Fig. S5A**) and increased VAV1 protein expression (**Fig. S5B**). These two mechanisms could account for resistance in 75% of the spontaneously resistant clones and populations (9 of 12). The remaining three clones (c2.2.1, c2.2.2, c7.1) all show increased phosphorylation of MEK1 on serine 298, indicating increased Pak activity in these cells (**Fig. S5B**). This observation suggests that these populations have active Rac/Cdc42 signaling, consistent with the DBL GEF-driven mechanism we identified.

We also evaluated the relevancy of our screen results with sequencing data from melanoma patients who had progressed on MAPKi treatment (31). Analysis of this previously published dataset of 18 patients (GSE65186) revealed that over 75% of patients (14 of 18) had over-expression of at least one resistance gene in the initial diagnostic biopsy or had acquired over-expression in at least one progression sample (**Fig. 4**). This suggests that melanomas with over-expression of resistance drivers at the time of treatment are less likely to respond to MAPKi and that acquired over-expression of our candidates during treatment may drive progression. However, analysis of a larger cohort is needed to determine of these trends are significant.

The identification of DBL GEFs as drivers of vemurafenib resistance also provides a direct mechanistic link between Src and vemurafenib resistance. Prior studies have implicated Src in mediating vemurafenib resistance (35, 39-41), but none have tied Src mechanistically to a pathway known to drive vemurafenib resistance in human melanoma. We have shown here that vemurafenib resistance driven by MCF2 and VAV1 can be blocked using the Src family kinase inhibitor saracatinib (**Fig. 5A**). Thus, one mechanism by which Src can drive vemurafenib resistance is through the activation of Rac/Cdc42/Pak through DBL GEFs such as MCF2 and VAV1. Importantly, we show that saracatinib blocks vemurafenib resistance driven by both over-expression of DBL GEFs and by over-expression of RAC1^P29S^, a previously identified mutation associated with vemurafenib resistance in cutaneous melanoma (24). Furthermore, we show for the first time that saracatinib can prevent the emergence of spontaneous vemurafenib resistance in long-term cultures of A375 cells (**Fig. 5D**).

Prior work has shown that N-terminal truncation of either BRAF^V600E^ or RAF1 promotes the formation of Ras-independent Raf dimers that cannot be inhibited with vemurafenib or dabrafenib(6, 37, 38). However, next generation Raf inhibitors have been developed that can block both monomeric and dimeric Raf(38). Importantly, one of these compounds, LY3009120, has recently been tested in a phase I clinical trial (NCT02014116). As predicted, we show that LY3009120 is more effective than vemurafenib at inhibiting the proliferation of melanoma cells expressing truncated BRAF^V600E^ or RAF1 (**Fig. 5F**). It is important to note, however, that LY3009120 was unable to completely inhibit the growth of these cells at the concentration used in our experiment. Nevertheless, the addition of saracatinib to LY3009120 was able to block growth of cells driven by all mechanisms we validated from our forward genetic screen (**Fig. 5F, 6A**).

We also evaluated a panel of MAPK inhibitors for their ability to control the growth driven by the various resistance drivers (**Fig. 6A**). As expected, each driver was able to support cell growth in the presence of either RAF inhibitors or MEK inhibitors alone or in combination. Interestingly, an ERK inhibitor was able to control growth driven by truncated BRAF or RAF1 but could not completely inhibit the growth driven by the DBL GEFs. This suggests that the DBL-driven drug resistance mechanism may involve a mechanism independent of MAPK. Nevertheless, DBL-driven resistance could be controlled with the addition of saracatinib.

Based on our findings, we propose a model of BRAFi resistance that involves two distinct mechanisms: one utilizing RAF/BRAF^V600E^ truncation and a second involving Rac1 activation via the DBL family members MCF2 and VAV1 (**Fig. 7**). Some aspects of the model that require additional studies. For instance, we do not yet understand the connection between BRAFi treatment and Src signaling. It is possible that MAPK inhibition leads to changes in Src activity through post-translational modification of the Src kinases and/or through transcriptional feedback mechanisms. It is also important to acknowledge that not all melanoma cell lines are responsive to saracatinib treatment, suggesting that not all melanomas show this BRAFi-induced Src alteration. Consistent with this idea, the DBL GEFs are unable to drive BRAFi resistance in some melanoma cell lines.

**Figure 7.**
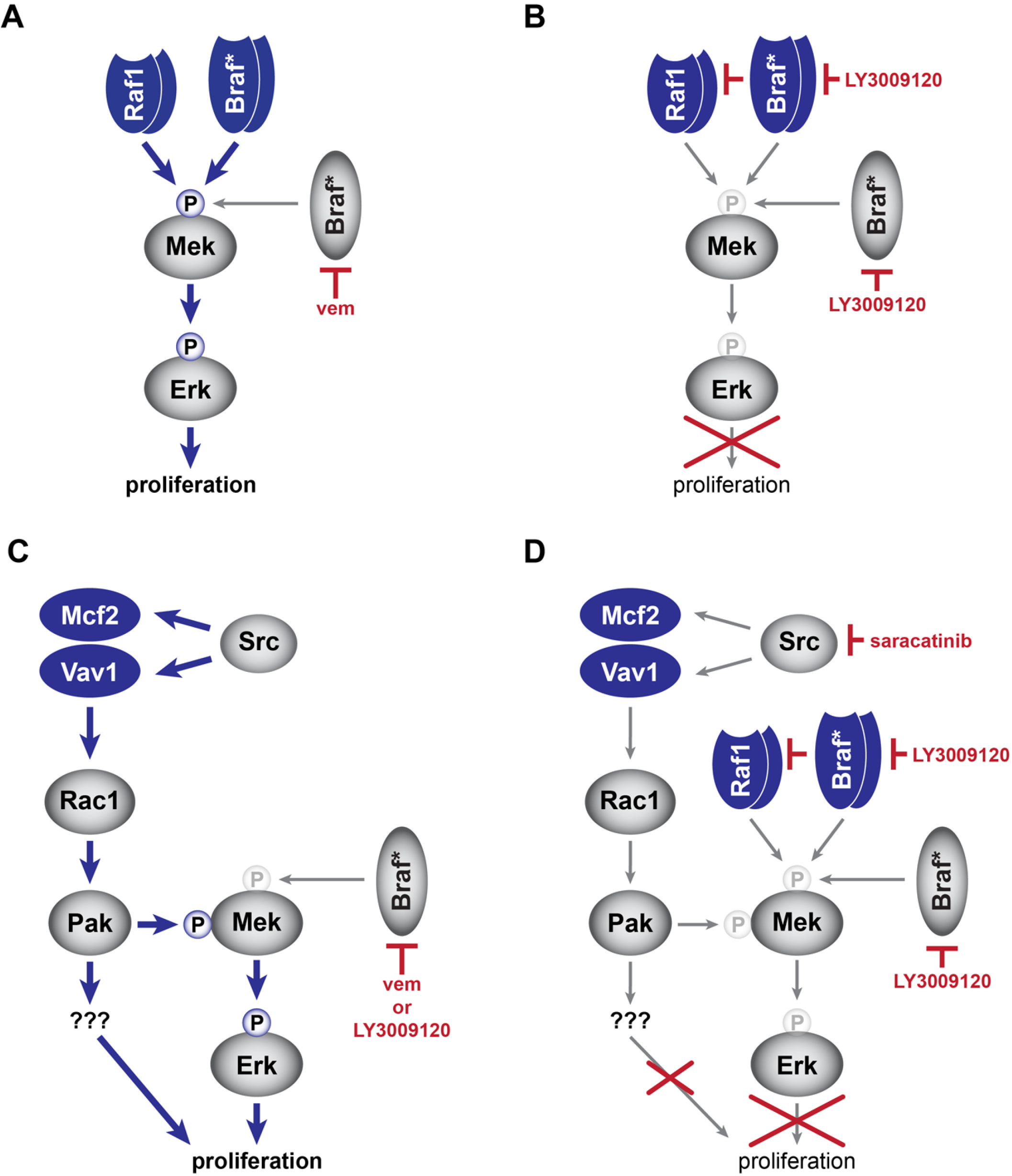
Model of drug resistance mechanisms. We have shown that N-terminal truncation of either RAF1 or BRAF^V600E^ leads to vemurafenib resistance (**A**) that can be overcome with the next generation RAF inhibitor LY3009120 (**B**). Our genetic screen identified a DBL-Rac1-Pak resistance mechanism that can drive proliferation in the presence of either vemurafenib or LY3009120 (**C**). However, the combination of saracatinib and LY3009120 can block both mechanisms identified by our screen (**D**).

Many of the questions raised by our experiments can be addressed by employing our forward genetic screening method in independent melanoma cell lines to elucidate shared and unique drivers of MAPKi resistance. Beyond resistance to vemurafenib, we can easily utilize our approach to identify novel mechanisms of resistance to next-generation inhibitors, such as LY3009120(37, 42). Lastly, our screening approach could inform the effective utilization of other targeted agents in cancer.

## Methods

### Sleeping Beauty Mutagenesis Screen

A375 cells were stability transfected via Effectene (Qiagen) coupled with a piggyBac transposase integration system (43) with Ef1α-SB100 transgene. After puro selection, SB100-expressing cells were transfected with the pT2/Onc3 transposon plasmid (21). 48 hours later, 1×10^6^ cells of SB100 + T2/Onc3 were plated on 10cm plates. Cells were subsequently treated with vemurafenib (5 µm), vemurafenib (5 µm) and cobimetinib (5 nm), or vehicle (DMSO, 0.2%) 24-hours after plating. Drug or vehicle was renewed every 3 to 4 days. Upon confluency (approximately 3-days after plating), the vehicle plates were collected. Cells treated with vemurafenib or vemurafenib with cobimetinib were collected after ∼18 or ∼28 days respectively.

To determine common transposon insertion sites across plates of resistant cells, genomic DNA from each plate was extracted using the GenElute^TM^ Mammalian Genome DNA miniprep Kit (Sigma). DNA fragments containing transposon/genome junctions were amplified via ligation mediated PCR and sequenced using the Illumina Hi-Seq 4000 platform as previously described (22).

### Colony Staining

1×10^6^ cells of either A375 SB100 + EGFP or SB100 + T2/Onc3 cells were stained with Coomossie Brilliant Blue after ethanol fixation 25 days after drug treatment (see screen details for concentrations). Colonies were counted with the GelCount^TM^ Colony Counter (Oxford Optronix).

### Creation of candidate overexpression constructs

Gene products mimicking the splice form driven by the SB transposon promoter were amplified from A375 cDNA (BRAF, RAF1, RAC1) or from a Transomics Technology human cDNA clones (MCF2, VAV1). Cloned cDNAs were inserted into piggyBac expression vector containing a human Ef1α promoter along with an IRES-puromycin-polyA cassette. Stable cell lines were obtained by co-transfection of each vector with a piggyBac transposase expression vector via Effectene (Qiagen) transfection reagent. Over-expression was assessed via RT-qPCR and immunoblot. See list of primers and antibodies for specifics.

### RNA Interference

RhoA and RhoC constructs had a pLKO backbone (Sigma-Aldrich). RAC1 constructs had a pZIP-mCMV vector backbone (Transomics Technologies). A non-targeting shRNA in the appropriate vector backbone was included to produce vector control cell lines. Cells were maintained as stably transduced, polyclonal populations. See Supplementary Information for RNAi targeting sequences.

### Cell culture

All cell lines were grown in DMEM (Gibco) supplemented with penicillin/streptavidin (Gibco) and 10% FBS (Gibco). Spontaneous vemerafinib-resistant clones and populations were created after 2-3 and 4-6 weeks cultured in 3 µm vemurafenib, respectively. Clones were isolated via cloning rings. All spontaneous resistant clones and populations were maintained in media with 3 µm vemurafenib.

### CellTiter Blue Viability Assay

Cells were plated in triplicate in 96-well plates at a density of 5 × 10^2^ to 5 × 10^3^ cell per well depending on the cell line. The CellTiter-Blue Viability Assay (Promega) was performed serially on pre- and post-inhibitor treated cells. Media containing the specific inhibitor used was renewed every 2-6 days. Fold change from Day 0 was assessed for each well by comparing pre- and post-inhibitor treated cells.

### Inhibitors

Inhibitors and the concentration used in these experiments include vemurafenib (5 µm; Selleckchem), cobimetinib (5 nm; Selleckchem), saracatinib (1 or 2 µm; Selleckchem;), FRAX486 (50 nm; Selleckchem), Fasudil (10 µm; Sigma), LY2009120 (Selleckchem; 1 µm), Imatinib (2 µm; Cayman Chemical).

### Immunoblotting

Rho protein activation assays were performed using sub-confluent 10 cm plates that were treated with the specified inhibitors for 48-hours prior to pulldown. Cells were lysed with ice cold 1% Triton X-100, 0.1% SDS, 10 mM MgCl2 with protease inhibitors, and activated Rho protein was recovered by pulldown with GST-PAK-CRIB fusion protein (44), followed by immunoblotting for active and total RAC1, CDC42, RHOQ, or RHOJ. Protocol adapted from Pellegrin and Mellor (45).

Antibodies used for immunoblotting were as follows: Rac1 (#610651, BD Transduction), CDC42 (#610928, BD Transduction), TC10 (RhoQ) ([Y304]ab32079, abcam), RhoJ (PA5-48271, Invitrogen), VAV1 (HPA001864, SigmaAldrich), p44/42 MAPK (ERK1/2) (#9102, Cell Signaling Technologies), phospho-p44/42 MAPK (T202/Y204) (#9101, Cell Signaling Technologies), MEK1 (#2352, Cell Signaling Technologies), phospho-MEK1 (Ser298) (#98195, Cell Signaling Technologies), phospho-AKT (S473) (#4060, Cell Signaling Technologies), AKT (610860, BD Transduction), β-actin (6221, BioLegend; A1978, Sigma), α-tubulin (12G10, DSHB). Secondary antibodies were IR antibodies from LiCOR or Rockland Inc. Immunoblots were imaged on a LiCOR Odyssey blot imager.

### Statistics

Statistical methods for some experiments are described within the text. The identification of transposon induced driver mutations was carried out using a gene-centric common insertion site method previously described (23).

