## Supplemental figures for "Src-dependent DBL family members drive resistance to vemurafenib in human melanoma"

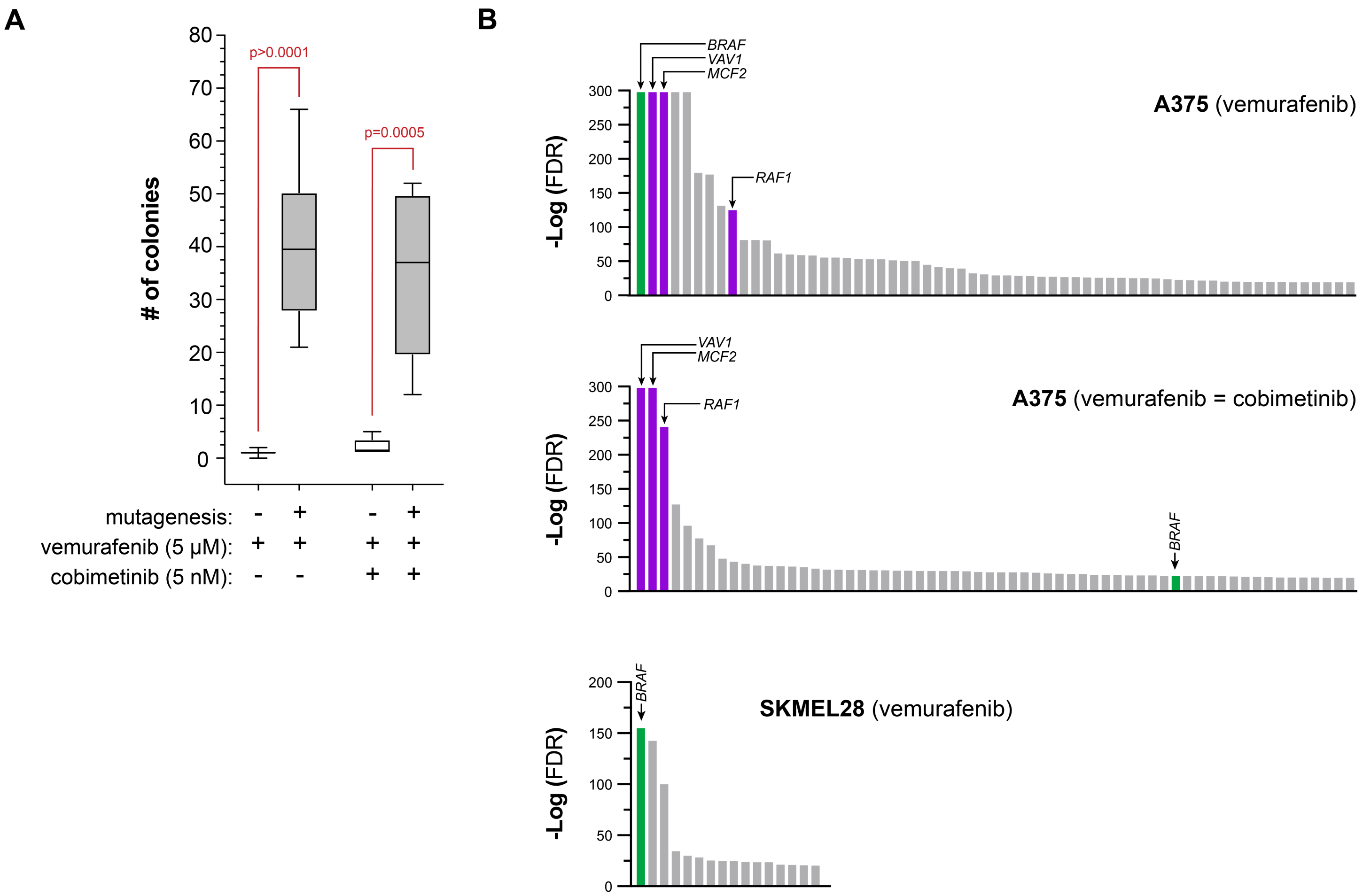


**Figure S1.** (**A**) Sleeping Beauty mutagenesis significantly increases the frequency of drug resistant colonies in A375 cells. (**B**) Plot showing significance [-log(FDR)] for genes identified in the indicated resistance screen.

[Green bar = significant gene identified in all screens, purple bar = significant gene identified in two screens, gray bar = unique to each screen].

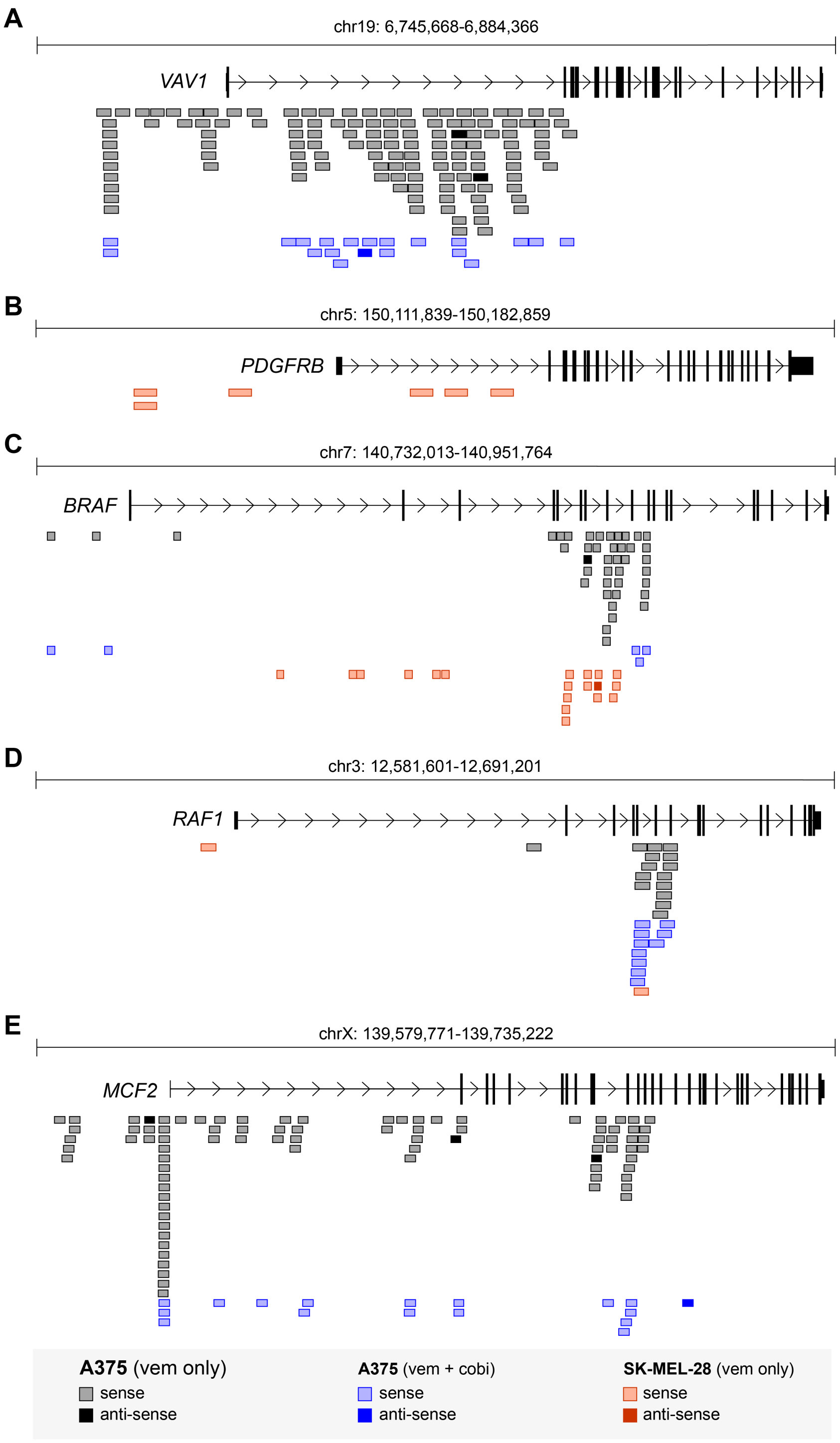


**Figure S2.** Distribution of transposon insertion events observed in the (**A**) VAV1, (**B**) PDGFRB (**C**) BRAF, (**D**) RAF1, and (**E**) MCF2. In all cases, rectangles represent the position and orientation of transposon insertions identified in the various screens (see legend).

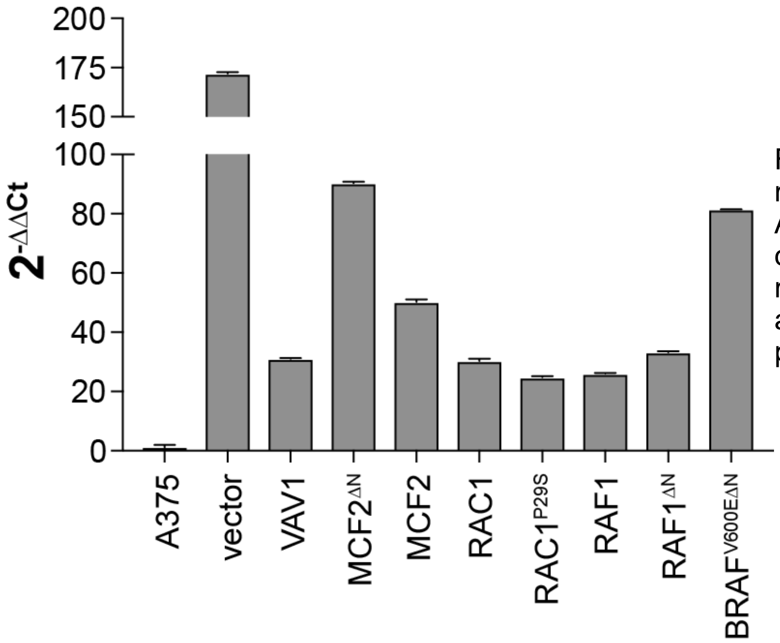


**Figure S3.** Quantitative RT-PCR analysis shows relative expression of each candidate gene in A375 cells. The relative expression values were obtained using primers to the puromycin resistance marker common to all constructs to allow for direct comparison across the various cell populations.

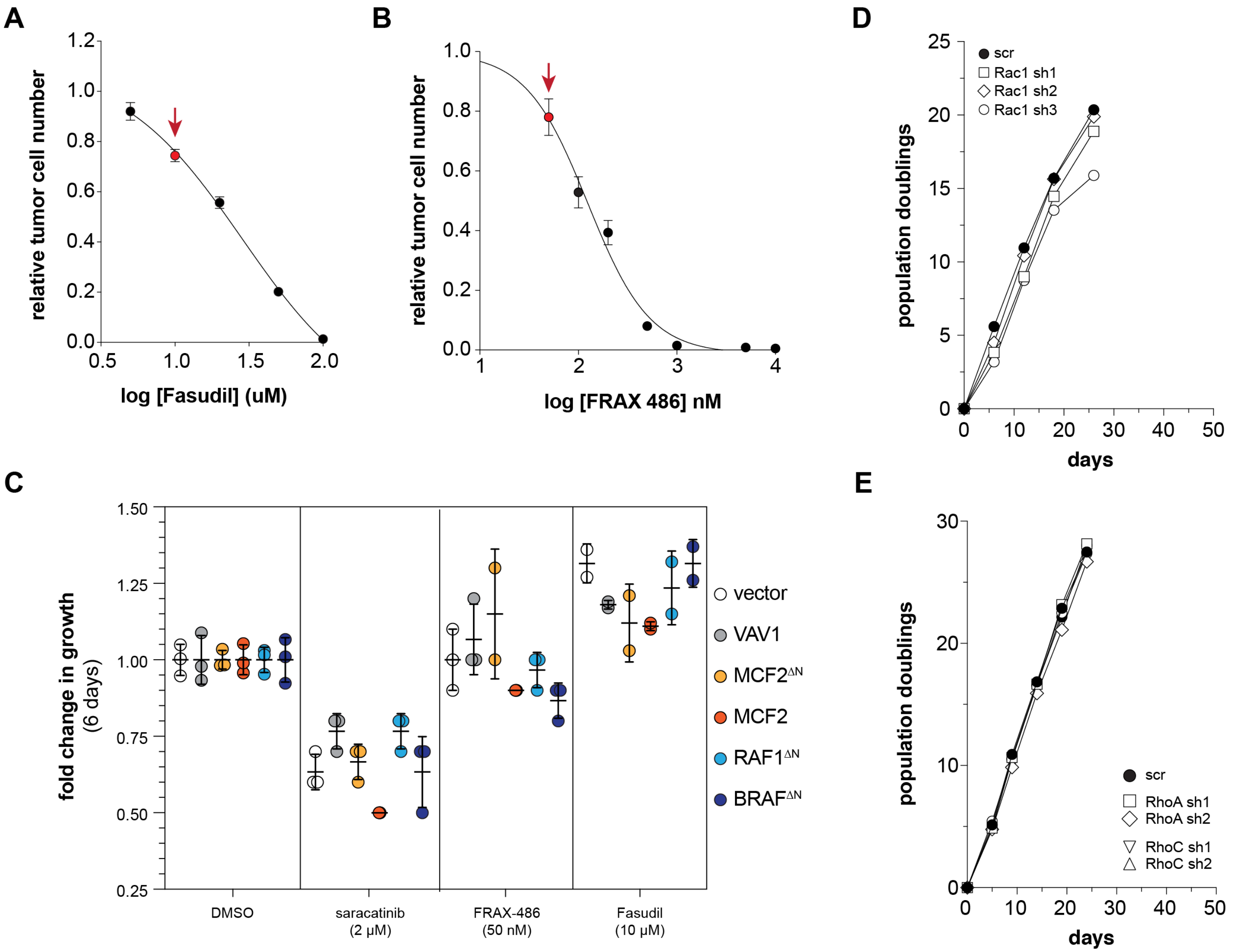


**Figure S4.** Characterization of drug and shRNA reagents used to interrogate the mechanism downstream of the DBL GEFs. A dose response curve was generated for Fasudi (**A**) and FRAX-486 (**B**) to identify a drug concentration that is well tolerated by A375 cells. [red dot = selected dose] (**C**) Cell growth assays using each drug at the selected concentration shows either no significant effect or slight growth reduction over a 6 day assay. Knockdown of Rac1 (**D**), RhoA, and RhoC (**E**) does not impact growth of A375 under standard culture conditions.

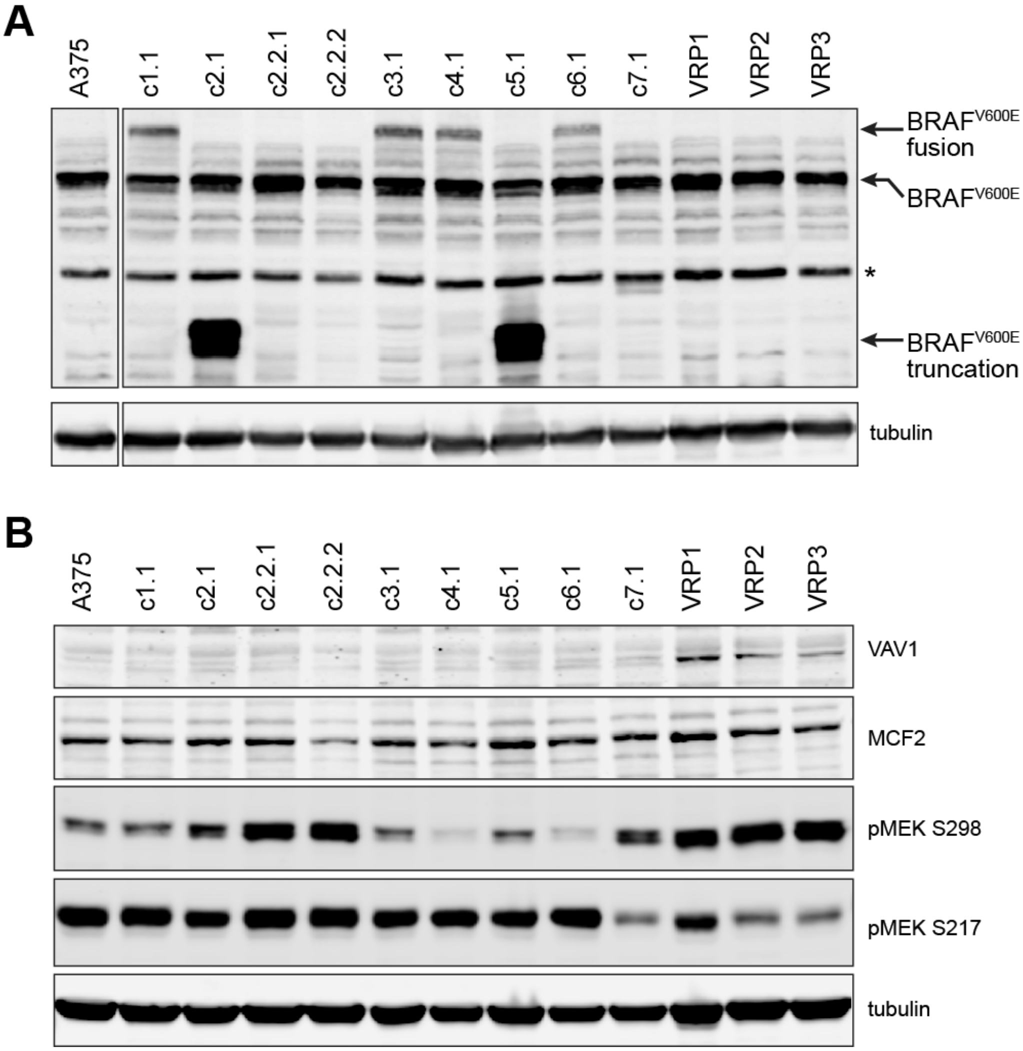


**Figure S5. Characterization of signaling in A375 clones that spontaneously acquired vemurafenib resistance.** (**a**) Western blotting using an anti-BRAF$^{V600E}$ antibody shows frequent alterations of BRAF$^{V600E}$ protein in spontaneously resistant A375 populations compared to parental A375 cells [*denotes a non-specific band]. (**b**) While MCF2 protein levels do not appear to change in resistant populations, VAV1 protein levels do appear to be increased in three independent cell populations. MEK phosphorylation on S217 (Raf site) and S298 (Pak site) also appear to be variable across the populations.
